# Interspecific introgression mediates adaptation to whole genome duplication

**DOI:** 10.1101/636019

**Authors:** Sarah Marburger, Patrick Monnahan, Paul J. Seear, Simon H. Martin, Jordan Koch, Pirita Paajanen, Magdalena Bohutínská, James D. Higgins, Roswitha Schmickl, Levi Yant

## Abstract

Adaptive gene flow is a consequential evolutionary phenomenon across all kingdoms of life. While recognition of widespread gene flow is growing, examples lack of bidirectional gene flow mediating adaptations at specific loci that cooperatively manage core cellular processes. We previously described concerted molecular changes among physically interacting members of the meiotic machinery controlling crossover number and distribution upon adaptation to whole genome duplication (WGD) in *Arabidopsis arenosa*. Here we conduct a population genomic study to test the hypothesis that escape from extinction following the trauma of WGD was mediated by adaptive gene flow between *A. arenosa* and its congener *Arabidopsis lyrata*. We show that *A. lyrata* underwent WGD more recently than *A. arenosa*, indicating that specific pre-adapted alleles donated by *A. arenosa* underwent selection and rescued the nascent *A. lyrata* tetraploids from early extinction. At the same time, we detect specific signals of gene flow in the opposite direction at other functionally interacting gene coding loci that display dramatic signatures of selective sweep in both tetraploid species. Cytological analysis shows that A. lyrata tetraploids exhibit similar levels of meiotic stability as *A. arenosa* tetraploids. Taken together, these data indicate that bidirectional gene flow allowed for an escape from extinction of the young autopolyploids, especially the rare tetraploid *A. lyrata*, and suggest that the merger of these species is greater than the sum of their parts.

## Introduction

Whole genome duplication (WGD) and hybridisation are key drivers of genomic novelty, promoting diversification in all kingdoms of life^1-3^. Recent progress in evolutionary genomics underscores the ubiquity of WGD at both ancient and recent time scales^4^, and population genomic approaches reveal widespread evidence of gene flow between the most diverse of species^5,6^. Both processes have therefore been associated with adaptive benefits. However, WGD and hybridisation are dramatic mutations, often leading to a host of genomic instabilities, including epigenetic shock, perturbed gene expression patterns, and meiotic instability, with direct negative impacts on fertility. Perhaps the most challenging issue is the most immediate: that of stable meiotic chromosome segregation following WGD. How nascent polyploids establish meiotic stabilisation remains an open unresolved question.

The wild outcrossing members of the *Arabidopsis* genus have recently emerged as fruitful models for the study of genome stabilisation following WGD^7^. *Arabidopsis arenosa* is a largely biennial outcrossing relative of the model *Arabidopsis thaliana*, which forms distinct lineages of diploids and autotetraploids throughout Central Europe^8-11^. Initial resequencing of a handful of autotetraploid *A. arenosa* individuals suggested selective sweep signatures at genes involved in genome maintenance, including DNA repair, recombination and meiosis^12^. Later, a targeted resequencing effort focused on patterns of differentiation between diploid and autotetraploid *A. arenosa*, revealing evidence of highly localised selective sweep signatures directly overlapping eight loci that interact during prophase I of meiosis^13^. These eight loci physically and functionally interact to control crossover designation and interference, strongly implying that a modulation of crossover distribution was crucial for polyploid establishment in *A. arenosa*^14,15^. Cytological evidence of a reduction in crossover numbers in the autotetraploids indicated that the selected alleles had an effect^13^. Similar to its sister species *A. arenosa* (*arenosa* hereafter), *Arabidopsis lyrata* (*lyrata* hereafter) also naturally occurs as diploids and tetraploids in well-differentiated lineages in central Austria^8,16-18^. While there is little evidence for gene flow among diploids of each species, there have been reports of gene flow between tetraploid *arenosa* and *lyrata* and, less pronounced gene flow between diploids and tetraploids^8,19,20^.

Here, we investigate the molecular basis of parallel adaptation to WGD in *lyrata* compared to *arenosa* and the possibility of adaptive gene flow between the two species. Specifically, we asked (1) whether the same or different loci may be involved in adaptation to WGD in *lyrata* as we observed in *arenosa*, and (2) whether these adaptations arose independently or via introgression from one species into the other. Using whole genome sequence data from 92 individuals of *lyrata, arenosa* and outgroups *Arabidopsis croatica* and *Arabidopsis halleri*, we first analysed population structure and demography, concentrating on assessing admixture and the degree and timing of population divergences. Then, to estimate the relative degree of adaptation to WGD across the ranges of *lyrata* and *arenosa*, we cytologically assessed meiotic stability in key populations. Finally, after scanning the *lyrata* genomes for signatures of selective sweeps, we compared outliers to those we previously found in *arenosa*^12,13^ and tested whether these selective sweep signatures overlap with fine-scale conspicuous introgression signals. Overall, our results reveal the molecular basis by which WGD was stabilised in both species and indicate that WGD-facilitated hybridisation allowed for stabilisation of meiosis in nascent autotetraploids by highly specific, bidirectional adaptive gene flow.

## Results & Discussion

### Population structure, demographic analysis and broad-scale admixture patterns

To understand population and species relationships we analysed the genomes of 92 individuals from ∼30 populations of *lyrata* and *arenosa* throughout Central Europe along with outgroups, sequenced at a depth of ∼15x per individual (*SI Appendix*, Table S1; Figure S1). STRUCTURE and Principal Component Analysis (PCA) showed a clear species-specific clustering for diploids, while tetraploids exhibited a gradient of relatedness between species (Figure 1A, B). Admixture was markedly lower in *arenosa* populations than in *lyrata*: consistently all diploids tested (SNO, KZL, SZI, BEL) as well as tetraploids from the Western Carpathians (TRE) and most of the Alpine tetraploids (HOC, GUL, BGS) harboured essentially pure *arenosa* genomes (Figure 1A). Minimal admixture signal (less than 1%) with *arenosa* was detected in a few *lyrata* genomes, in particular the Austrian diploid (PEQ, PER, VLH) as well as the *lyrata* eastern tetraploid (*Let* hereafter) populations (LIC, MOD), and the tetraploid KAG population (Figure 1A).

**Figure 1.**
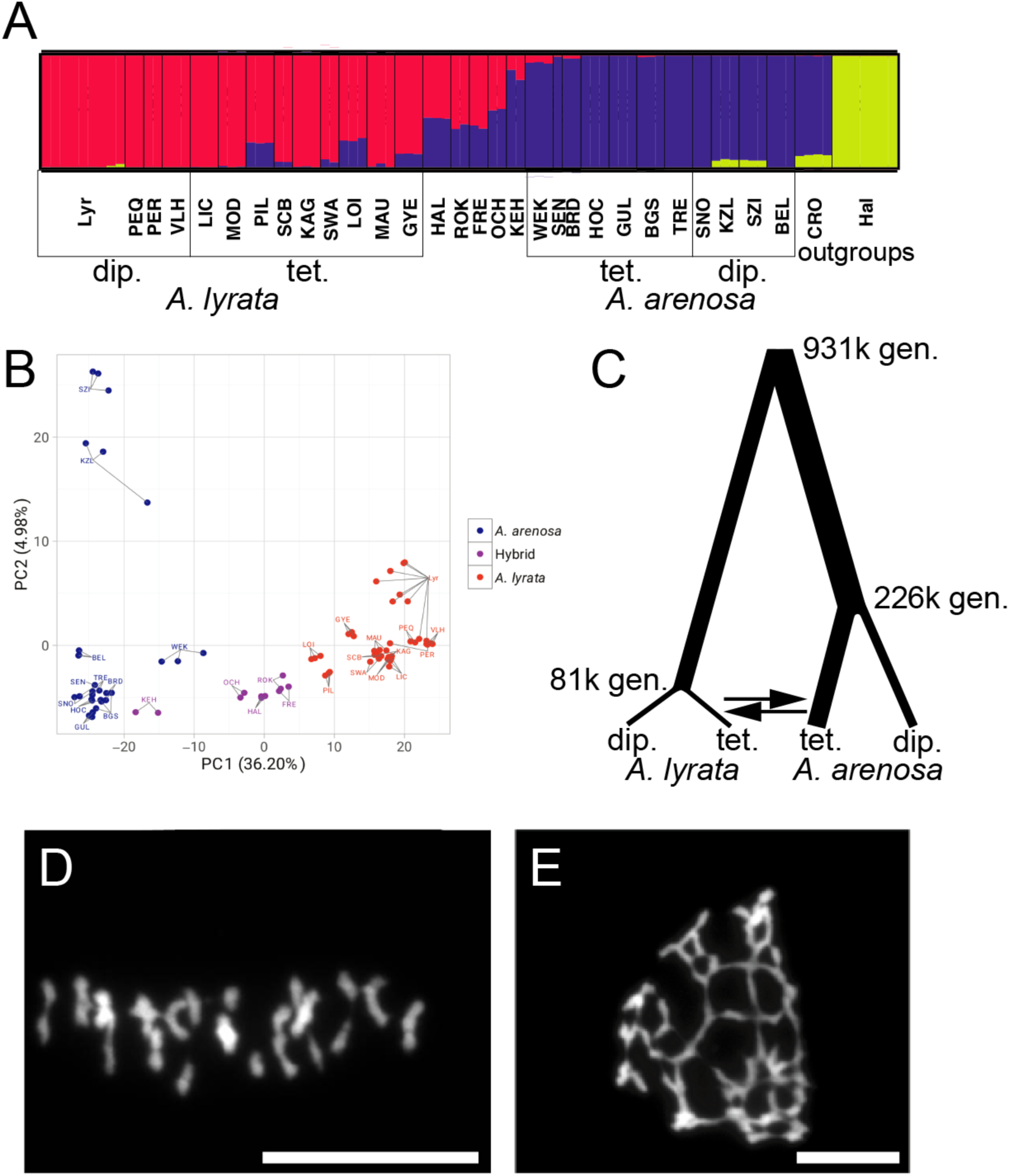
Ploidy-specific admixture, demographic history, and stable autotetraploid meiosis in *A. lyrata*. (A) A continuous range of admixture specifically in tetraploid populations demonstrated with STRUCTURE analysis of nuclear SNP data (32,256 LD-pruned, 4-fold degenerate SNPs). Populations (in three letter code) and population groupings (ploidy, species) designations are displayed. (B) PCA shows individuals group on the main (PC1) axis by species and not by ploidy, with hybrid individuals located between *A. lyrata* and *A. arenosa* samples. We call hybrids all non-pure populations from the hybrid zone in the eastern Austrian Forealps (see *SI Appendix*, Figure S1). (C) Demographic parameter estimates for *A. lyrata* and *A. arenosa* populations. Line widths are proportional to estimates given in *SI Appendix*, Figure S2. (D and E) Meiotic chromosome spreads of tetraploid *A. lyrata* during metaphase I: (D) Stable meiosis, with rod and ring bivalents exhibiting proximal and distal chiasmata, respectively. (E) Unstable meiosis, with multivalents and multiple chiasmata on chromosome arms. Chromosomes are stained with DAPI; bar = 10 µm.

In contrast, many populations exhibited substantial admixture signals between *lyrata* and *arenosa*, contrasting drastically in degree (Figure 1A). Several tetraploid *lyrata* populations from the Wachau (SCB, SWA, MAU) displayed only slight admixture with *arenosa*, and populations at the Wachau margin (PIL, LOI) showed stronger admixture, probably due to the increased proximity to the Hercynian and Alpine *arenosa* lineages^21^. Compared to the Wachau, where *lyrata* occurs on the slopes and hilltops along the Danube river surrounded by *arenosa* populations outside of the valley, there is a classical hybrid zone in the eastern Austrian Forealps: the parental species are found at the two poles of the zone (diploid and tetraploid *lyrata* in the Wienerwald, tetraploid *arenosa* at higher altitudes to the west, the hybrids between, Figure S1). *Lyrata* populations HAL, ROK, FRE, OCH and KEH are heavily admixed with *arenosa*, with KEH appearing more *arenosa*-like and FRE being more *lyrata*-like; HAL, ROK and OCH showed intermediate genetic clusters (Figure 1A). Again, proximity of the hybrids to the donor species corresponded with increased admixture. The Hungarian tetraploid *lyrata* population GYE also exhibited admixture signal, suggesting that gene flow between *lyrata* and *arenosa* is not restricted to the Austrian Forealps. Principle Component Analysis was consistent with STRUCTURE findings, with PC1 dividing samples by species (explaining >36% of the variance; Figure 1B). The second axis (<5% of the variance) separated KZL and SZI from the other diploid *arenosa* populations. These are representatives of the Pannonian lineage, which is the oldest and most distinct diploid *arenosa* lineage^21^. Overall, our results are consistent with previous descriptions of introgression between *lyrata* and *arenosa* in Austria that were based on smaller marker sets and different sampling schemes^8,22^.

We estimated the population split time between *lyrata* and *arenosa* at 931,000 (931k) generations using *fastsimcoal2*^23^ (Figure 1C; *SI Appendix*, Figures S2, S3, Table S2). This translates to ∼2 million years (my), given an average generation time of 2 years, which would coincide with the onset of Pleistocene climate oscillations. This estimate lies within the range of age estimates for this split from ^24^ with 1.3my and ^25^ with 8.2my. We estimated the age of WGDs at 81k generations for *lyrata* (∼160,000 years) and 226k generations for *arenosa* (∼450,000 years), which approximately mark periods of glacial maxima^26^. Noting this, we next asked if either species experienced substantial historical bottlenecks. Using Pairwise Sequentially Markovian Coalescent Model (PSMC^27^) we could not reach ages as ancient as 130-300kya (*SI Appendix*, Figure S4) because the recombinant blocks that PSMC measures are too short in these diverse outcrossing species to estimate ancient population histories. Our analysis did reveal that diploid *lyrata* had a peak effective population size (*Ne*) ∼25kya (PER, VHL) and ∼20kya (PEQ), while diploid Dinaric *arenosa* (BEL) peaked earlier, at ∼30kya. Interestingly, diploid Western Carpathian *arenosa* SNO, the population that founded several widespread autotetraploid lineages^9^, gave a strong signal of continuous expansion. These results suggest that diploid *lyrata* and *arenosa* underwent a bottleneck after the last glacial maximum 30-19kya. PSMC does not accommodate autotetraploid data, but using *fastsimcoal2*, we detected a strong bottleneck at WGD for *lyrata*, but none for *arenosa* (*SI Appendix*, Figure S2).

We next assayed for patterns of gene flow using coalescent modelling with *fastsimcoal2*. Due to model overfitting when using more than two migration edges, we chose the model retaining only two migration edges with highest support: interspecific gene flow from tetraploid *arenosa* to tetraploid *lyrata* (0.0998 alleles/generation) and *lyrata* to *arenosa* gene at the same level (0.1001 alleles/generation) (Figure 1C; *SI Appendix*, Figure S2, S3, Table S2).

To identify particular genomic regions harbouring signatures of gene flow, we first used *D*_FOIL_^28^. In total, we identified 127 genomic windows across all scenarios suggesting introgression between tetraploid *lyrata* and *arenosa* (*SI Appendix*, Figure S5). Across all tested scenarios, the most prevalent introgression signal was ancestral (i.e. prior to the diversification of the *lyrata* tetraploids). This fits well with the recent suggestion that introgression between *lyrata* and *arenosa* began during the penultimate interglacial^22^. It should be noted that the directionality of ancestral introgression cannot be determined by *D*FOIL. In case of more recent introgression, our results indicate that directionality was primarily from *arenosa* into *lyrata* (*SI Appendix*, Figure S5). This supports greater levels of gene flow with *lyrata* serving as plastid donor, at least recently (consistent with earlier findings^8^). These findings partly contrast with^20^, who found that crosses between *lyrata* and *arenosa* tetraploid gave viable seeds in both directions, but that the cross with maternal *arenosa* performed better. We note that each study used different populations, which may explain contrasting results in these diverse outcrossing species. We note also that selection based on exogenous factors may currently favour a particular direction of gene flow^29^, and such selection might have been absent in the past and certainly in experimental crosses.

### Stabilisation of *lyrata* meiosis following WGD

Given the very low abundance of *lyrata* tetraploids in nature, we assayed whether these tetraploids were indeed meiotically stable. Cytological analysis indicated that in fact *lyrata* tetraploids exhibit similar levels of meiotic stability as *arenosa* tetraploids, as evidenced by relative percentages of stable rod and ring bivalents (Figure 1D; *SI Appendix*, Table S3) vs. less stable multivalents (Figure 1E; *SI Appendix*, Table S3). We were surprised to observe among both species that meiotic stability segregates within populations, typically ranging from <20 to >60% stable metaphase I cells per plant with extremes observed in LIC (0 to 100%) and consistently low levels (<20%) in the SEN population. Overall these results indicate that meiotic stability is broadly segregating within tetraploid populations of both species, as well as the hybrids.

### Selective sweep signatures at loci involved in adaptation to WGD in *lyrata*

To gain insight into the processes underlying adaptation to WGD in *lyrata* tetraploids, we performed a population-based genome scan for selection. We quantified differentiation between *lyrata* ploidies by calculating *dXY*, Fst, and Rho in adjacent windows along the genome between diploids and tetraploids. We focused on the non-admixed *lyrata* tetraploid populations LIC and MOD (*lyrata* eastern tetraploids; *Let*), which by STRUCTURE and PCA analyses exhibited the lowest levels of admixture (Figure 1A) and clustered with *lyrata* diploids, distant from the *arenosa* tetraploids or the broadly admixed *lyrata* tetraploids (Figure 1B). Overall, genome-wide differentiation between *lyrata* diploids and the tetraploid populations was extremely low (mean Rho between ploidies = 0.19) with few fixed differences (Table 1; *SI Appendix*, Table S4 for additional population contrasts), consistent with our previous studies in *arenosa*^12,13,21^.

**Table 1.** Genome-wide differentiation between *A. lyrata* diploids and tetraploids and between tetraploid lineages grouped by biogeography. Differentiation metrics shown are allele frequency difference (AFD), *dXY*, Fst, Rho, and the number of fixed differences (Fixed Diff).

| Contrast | No. SNPs | AFD | $d_{XY}$ | Fst | Rho | Fixed Diff |
| --- | --- | --- | --- | --- | --- | --- |
| Diploid *lyrata* vs. *lyrata* eastern tetraploids (*Let*) | 2,904,110 | 0.14 | 0.22 | 0.09 | 0.19 | 270 |
| Diploid *lyrata* vs. *lyrata* Wachau tetraploids (*Lwt*) | 3,794,257 | 0.11 | 0.16 | 0.07 | 0.17 | 64 |
| *Lyrata Let* tetraploid vs. *Lwt* tetraploid | 4,795,381 | 0.09 | 0.16 | 0.06 | 0.13 | 24 |
| *Arenosa* Hercynian tetraploids vs. *arenosa* Alpine tetraploids | 1,812,223 | 0.10 | 0.16 | 0.03 | 0.07 | 0 |

Contrasting the *lyrata* diploids and the *Let* tetraploids, we partitioned the genome into gene-sized windows (*SI Appendix* Methods), and identified the extreme 1% empirical outliers for allele frequency differences (AFD), *dXY*, Fst, Rho, and the number of fixed differences. When comparing the inclusive set of these outliers to the extreme 0.5% outliers reported for the diploid-autotetraploid contrast in *arenosa* in^13^, we observed selective sweep signatures at the majority of the known loci that were found as top outliers in *arenosa*, having primary functions of mediating meiosis, endopolyploidy, and transcription. In particular outlier loci participate in meiotic crossover formation, including *PRD3* (Figure 2), *ASY1, ASY3, PDS5b* and *SYN1*. These results indicate that the same processes were under selection following the more recent WGD event in *lyrata* as were under selection following the independent, earlier WGD (Figure 1C) in *arenosa*.

**Figure 2.**
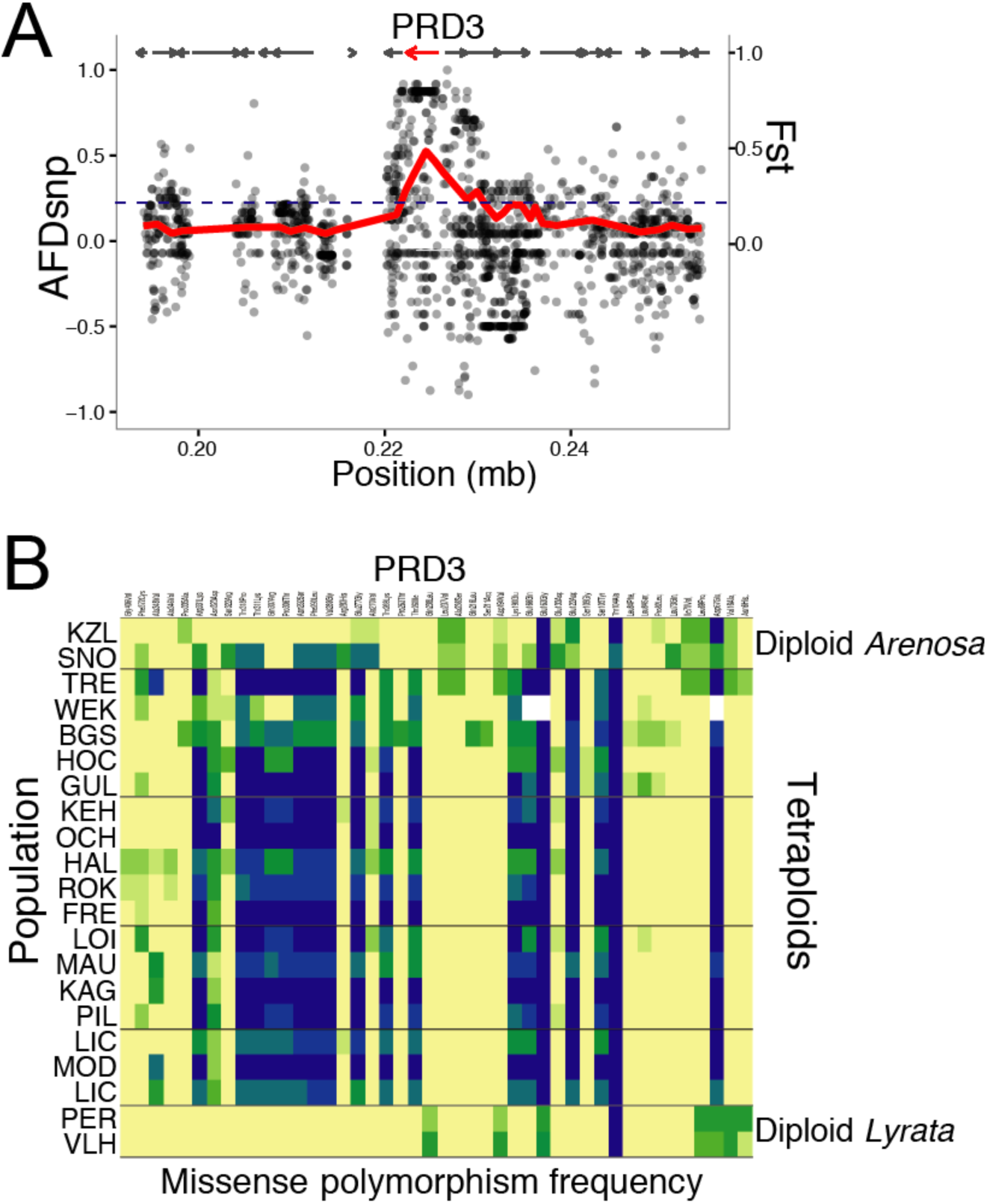
Selective sweeps and missense polymorphism frequencies by population. (A) *A. lyrata* selective sweep in *PRD3*, a gene involved in meiotic double strand break formation. Dots represent polymorphic SNPs. (B) Heatmap representing allele frequencies of missense polymorphisms in *PRD3*. Frequencies 0-100% follow yellow to green to blue. Derived diploid *A. arenosa*-specific missense polymorphisms are driven to high frequency in the tetraploids, while diploid *A. lyrata* alleles are absent, implicating diploid *A. arenosa* origin to this selected allele in the tetraploids.

Apart from loci encoding meiosis-related genes, we detected extreme differentiation at loci belonging to other functional categories clearly related to the challenges attendant to WGD, including loci involved in endoreduplication and transcriptional regulation: *CYCA2;3, PAB3, NAB, TFIIF*, and *GTE6*/*NRPA1* (Table 2).

**Table 2.** Overlap list of the top 1% outliers from the genome scan of diploid *A. lyrata* vs. *Let* (“*Let* scan”) and diploid *A. lyrata* vs. *Lwt* (“*Lwt* scan”), overlapped with the outliers of the *A. arenosa* diploid-tetraploid scan of ^13^. The overlap between the diploid/*Let* and diploid/*Lwt* contrasts yielded 196 genes, which is approximately a third of the gene identified in each scan. The overlap of those two scans with the *A. arenosa* scan gave 20 genes in common. Core meiosis genes found in Yant et al (2013) that were found in only one or none of the two *lyrata* scans are stated in the bottom part of this list.

| *Lyrata* ID | Name | Description | *Let* scan<br>Outlier | *Lwt* scan<br>Outlier |
| --- | --- | --- | --- | --- |
| *AL1G10680* | *PRD3* | Involved in meiotic double strand break formation. | Yes | Yes |
| *AL1G27690* | *CYCLIN A2;3* | A2-type cyclin. Negatively regulates endocycles and acts as a key regulator of ploidy levels in *Arabidopsis* endoreduplication. Interacts physically with CDKA;1. Expressed in trichomes and young tissues. | Yes | Yes |
| *AL1G36300* | *POLY(A) BINDING PROTEIN 3 (PAB3)* | Putative poly(A) binding protein. May therefore function in posttranscriptional regulation, including mRNA turnover and translational initiation. Expression only in floral organs. | Yes | Yes |
| *AL2G25520* | *SWEETIE* | Involved in trehalose metabolic process, carbohydrate and starch metabolic process | Yes | Yes |
| *AL2G25920* | *ASY1* | Meiotic asynaptic mutant 1. ASY1 protein is initially distributed as numerous foci throughout the chromatin. During early G2 foci appear on nascent chromosome axes to form an axis-associated signal. | Yes | Yes |
| *AL2G37810* | *PDS5-like* | ARM repeat superfamily protein. | Yes | Yes |
| *AL2G40680* | *CHROMO-METHYLASE 1 (CMT1)* | Ecotype KI-0 chromomethylase (CMT1). A plant line expressing an RNAi construct directed against DMT4 has reduced agrobacterium-mediated tumor formation. | Yes | Yes |
| *AL4G29630* | *NAB* | Nucleic acid-binding, OB-fold-like protein | Yes | Yes |
| *AL4G29650* | *unknown* | Unknown protein | Yes | Yes |
| *AL4G30770* | *MEE22* | Involved in endoreduplication and maintenance of meristem cell fate. | Yes | Yes |
| *AL4G46460* | *ASY3* | Coiled-coil domain protein required for meiosis. | Yes | Yes |
| *AL5G13440* | *ASF* | Asparagine synthase family protein | Yes | Yes |
| *AL5G32850* | *PSF* | Pseudouridine synthase family protein | Yes | Yes |
| *AL5G32860* | *TFIIF* | Transcription initiation factor IIF, beta subunit: Functions in RNA polymerase II transcription activity. | Yes | Yes |
| *AL5G32870* | *GTE6* | Bromodomain containing nuclear-localised protein involved in leaf development. | Yes | Yes |
| *AL5G39280* | *NRPA1* | A subunit of RNA polymerase I (aka Pol A). | Yes | Yes |
| *AL6G15380* | *SYN1* | Encodes a RAD21-like gene essential for meiosis. | Yes | Yes |
| *AL7G35790* | *unknown* | Unknown protein | Yes | Yes |
| *AL8G25590* | *DYAD, SWI1* | Involved in sister chromatid cohesion and meiotic chromosome organization in male and female meiosis. | Yes | Yes |
| *AL8G25600* | *TPR-like* | Tetratricopeptide repeat (TPR) protein | Yes | Yes |
| *AL1G35730* | *ZYP1a, b* | Transverse filament of SC | No | Yes |
| *AL4G20920* | *SMC3* | Member of *A. th.* meiotic cohesin complex | No | No |
| *AL8G10260* | *PDS5* | Member of *A. th.* meiotic cohesin complex | No | Yes |

Whole genome duplication increases the ploidy of all cells, while endopolyploidy occurs in single cells during their differentiation, and this cell- and tissue-specific ploidy variation is important in plant development^30-32^. Thus, given the instantly doubled organism-wide nuclear content following WGD, we postulate that the degree of endopolyploidy would be modulated in response, with accumulating support this notion^33-35^. Our findings bolster the idea that there may be a link between organism-wide polyploidization and that of single cells within an organism. Research about the effect of WGD-induced dosage responses of the transcriptome is still in its infancy^3,36^, and emerging studies on allopolyploids also support incomplete dosage compensation.

While the comparison of the most pure *lyrata* tetraploid populations, represented by the *Let* group, to *lyrata* diploids is the most stringent test of which loci are under selection in a purely *lyrata* genomic context, we extended our tetraploid *lyrata* sampling to populations from the Wachau, which frequently showed admixture with *arenosa* (*lyrata* Wachau tetraploids, *Lwt* hereafter: PIL, SCB, KAG, SWA, LOI, MAU). Because the *Let* and part of the *Lwt* populations grow in contrasting edaphic conditions (*Let* on limestone, *Lwt* on siliceous bedrock), we used this approach to maximise our chances of capturing differentiation specifically related to ploidy and not local adaptation. We observed that differentiation between these two tetraploid *lyrata* groups is stronger than differentiation between the tetraploid *arenosa* lineages studied here (Table 1), suggesting that there is stronger genetic structure within *lyrata* than *arenosa*, as was observed by ^37^. While the top 1% outlier genes were identified independently for the *Let* and *Lwt* (636 for the *Let*, 669 for the *Lwt*; *SI Appendix*, Dataset S1), we found considerable overlap (196 gene-coding loci, approximately a third of loci identified in each scan; *SI Appendix*, Dataset S1). Genes encoded by these loci include many involved in genome maintenance as well as meiosis. Comparison of this list with the outliers reported following WGD in *arenosa*^13^ yielded 20 gene-coding loci that exhibited the highest levels of differentiation in both studies (Table 2). These included those meiosis-related loci reported above for the *Let* contrast (*PRD3, ASY1, ASY3*, and *SYN1*), as well as the endopolyploidy genes *CYCA2;3* and *MEE22*, and the global transcriptional regulator *TFIIF*, among others. In addition, the important meiosis controlling loci *ZYP1b* and *PDS5* were outliers in the *Lwt* contrast. *ZYP1b* was moderately differentiated in the *Let* group also, but was not among the 1% top outliers; *PDS5* showed no differentiation between the *Let* and *lyrata* diploids. *SMC3*, a top outlier in *arenosa*, showed only moderate differentiation in the *Lwt* and no differentiation in the *Let*. These results further support the notion that these same loci governing meiotic recombination, endoreduplication, and transcription are important for adaptation to WGD and that signatures observed at these loci are not signatures of local adaptation. Further, it indicates that selected alleles are still segregating throughout the hybrid zone.

### Highly specific introgression and candidate loci for adaptive gene flow

Finally, we sought to confirm whether the strong observed signals of selective sweep were the products of interspecific introgression. To confirm candidate introgressed regions at high resolution we used *Twisst*^38^, performing two independent analyses, with either *Let* or *Lwt* representing tetraploid *lyrata*. The consensus species phylogeny, topology 3, indicated the overwhelmingly dominant genome-wide topology (Figure 3A, B).

**Figure 3.**
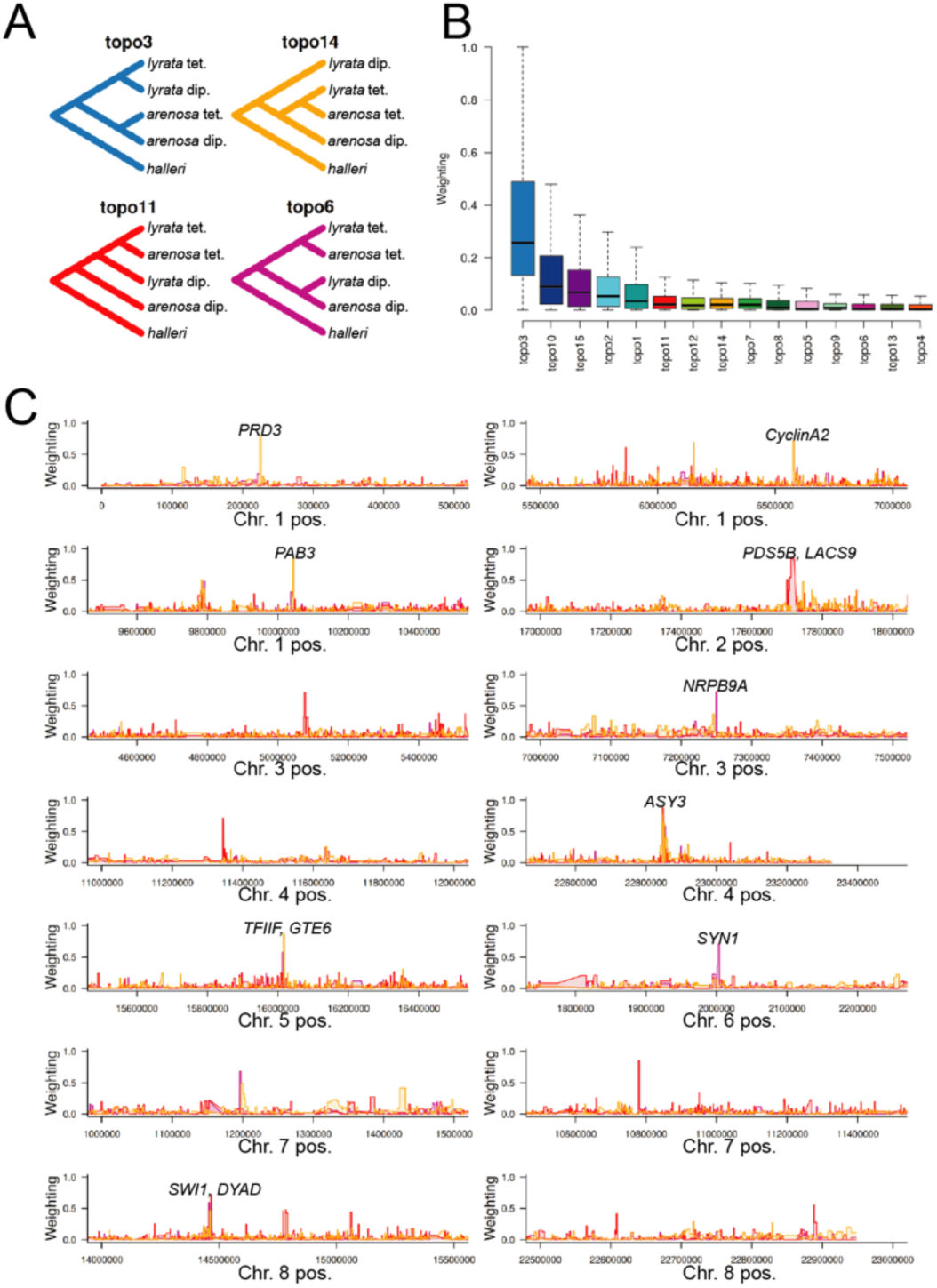
Highly specific introgression events across species boundaries. (A) Selected topologies from *Twisst* analysis of *Lwt*: topology 3 is the dominant species tree, topologies 11, 14 and 6 indicate localised gene flow between tetraploids. (B) Relative weightings of all topologies across the genomes analysed. (C) Introgression events revealed by *Twisst* analysis are highly localised at loci encoding genes controlling meiosis, endopolyploidy and transcriptional control. All gene coding loci under a given narrow peak are labelled; all indicated loci are divergence scan outliers in both the *Let* and *Lwt* divergence scans in addition to being Twisst outliers. The genome-wide dominant topology 3 weightings are omitted in panel C for clarity.

Of windows exhibiting high weightings (greater than 0.5) of the three introgression-indicative topologies (topology 3, 11, or 14) we found clear peaks overlapping many of our divergence scan outliers (Figure 3C; *SI Appendix*, Dataset S2). In total, 21 gene coding loci that were positive in both the *let* and *lwt* divergence scans also exhibited a weighting of 0.5 or greater for one of the introgression-indicative topologies (Figure 3C; *SI Appendix*, Dataset S2). This degree of overlap of the loci found under selection in our genome scans is dramatically greater than expected by chance, which we confirmed by performing permutation tests (*SI Appendix*, Figure S6). Like the divergence outlier windows, *Twisst*-positive windows were narrow, which might be an indication that genomic differentiation following divergence between *lyrata* and *arenosa* is advanced, similar to the numerous narrow genomic regions of introgression in the case of gene flow between *Populus alba* and *Populus tremula*^39^. From the *Twisst* analysis a slight majority of these discrete introgressed sweep loci appear to have a *lyrata* origin (Figure 3A). Topology 11 (direction: *lyrata* into *arenosa*) was marginally the most frequent of the three introgression-indicative topologies, being dominant in 14(*Lwt*)/29(*Let*) windows overall. Topology 14 (*arenosa* into *lyrata*) was dominant 11(*Lwt*)/20(*Let*) times, and 7(*Lwt*)/8(*Let*) windows were dominant for topology 6 (direction unclear).

Taking these results together, we observe that several meiosis-related loci were outliers in all *Twisst* and divergence scans: *PRD3, ASY3, SYN1*, and *DYAD* (*SI Appendix*, Dataset S2), with only four not showing a signal in both *Twisst* analyses as well as both divergence scans: *ASY1, ZYP1a, ZYP1b*, and *PDS5*. Overall, however, the vast majority of these loci were indicated as undergoing specific gene flow in one of the tests, with a slight preponderance of the directionality of genome-wide introgression from *arenosa* to *lyrata* (*fastsimcoal* and *Twisst* analyses together indicated largely bidirectional gene flow, while *D*FOIL supports nearly unidirectional recent gene flow from *arenosa* to *lyrata*). Given that *arenosa* is the much older tetraploid (twice as ancient as *lyrata*; Figure 1C) and much more widespread, we hypothesise that arenosa-sourced alleles were under selection for stable polyploid meiosis longer, providing preadapted alleles to the nascent *lyrata* tetraploid. Introgression of optimised alleles from an older to a younger species has been indicated for high-altitude adaptive alleles from Denisovans and Tibetan *Homo sapiens*^40^. Taking *PRD3* as a clear example (Figure 2B), derived diploid *arenosa*-specific missense polymorphisms are driven to high frequency in the tetraploids of both species, while diploid *lyrata* alleles are absent, strongly implicating diploid *arenosa* origin.

Our findings suggest that introgression of particular alleles of meiosis-related genes might stabilise polyploid meiosis, with the less effective alleles of one species being replaced by introgression of alleles from orthologous loci in the other tetraploid. Introgression of alleles optimised for adaptation to WGD could be especially beneficial in hybrid zones such this one, which spans a climatic gradient from a warmer, Pannonian climate at its eastern margin to harsher conditions in the eastern Alps. Meiosis is a temperature-sensitive process^15^, and we hypothesise substantial levels of meiosis-related allele-environment associations with variable temperature. Allele-environment associations with climatic variables across a hybrid zone have been observed in spruce^41^.

## Conclusion

For these newly formed tetraploids WGD appears to be both a blessing and a curse. Although WGD appears to have opened up access to the allelic diversity of a sister species, as well as provided population genomic benefits^21^, it also presents new challenges to the establishment of optimal allelic combinations. Because the gene products at the meiotic loci under the most extreme selection across this hybrid zone functionally and physically interact, we expect that efficient evolved polyploid meiosis requires the harmonious interactions of multiple selected, introgressed alleles in concert. However, relatively high levels of residual masking of genetic load in autotetraploids^21,42^ will tend to extend the duration that deleterious alleles segregate in populations, with negative phenotypic consequences. This is consistent with our observation that polyploid meiosis exhibits wide degrees of within-population variability in stability across this hybrid zone (*SI Appendix*, Table S3). This observed diversity suggests that the optimal combination of meiosis alleles is yet segregating, which may also be the result of the recent age of these WGDs. Dedicated molecular investigation of whether the measured within-population meiotic stability is associated with particular allele combinations is the focus of ongoing functional analyses.

In this study, we investigated the population genetic basis of adaptation to WGD in congeners that hybridise only as tetraploids. We found that many of the same loci exhibit the most conspicuous signatures of selective sweep in *lyrata* following WGD that we observed in *arenosa*, and further, that the strongest signals of interspecific introgression occur precisely at many of these same loci. Using whole genome sequence data from 30 populations we probed complex population structure and patterns of gene flow. Interestingly, we observed cytologically that the degree of meiotic stability varied dramatically, even within populations of both species, suggesting that stability has not been completely established, or that other, perhaps epigenetic or environmental factors influence meiotic stability in still unknown ways. At the same time, populations exhibited admixture signals that contrast dramatically in degree, indicating a complex introgression landscape. We present evidence that the molecular basis by which WGD was stabilised in *lyrata* and *arenosa* is shared. Our data further suggest that WGD-facilitated hybridisation allowed for stabilisation of meiosis in nascent autotetraploids by specific, bidirectional adaptive gene flow, tightly overlapping loci known to be essential for processes that are impacted by WGD: meiotic stability, endopolyploidy, and transcription, and others. It is curious that the very process that rescues fitness in these species, hybridisation, is potentiated by the same phenomenon to which the resultant adaptive gene flow responds: WGD.

## Methods

### Sample design, genomic library preparation and sequencing

Individual plants were collected from field sites across Central Europe (*SI Appendix*, Figure S1). Cytotypes were determined by flow cytometry from these populations in ^8^ and ^21^; no triploids have been detected in these populations, nor have we found any evidence in the flow cytometry or cytology data that any of these populations consist of mixed-ploidy subpopulations.

Central European tetraploid *lyrata* has its largest distribution in eastern Austria, in two biogeographic regions: the Wienerwald (*lyrata* eastern tetraploids / *Let* hereafter; LIC, MOD), and the Wachau (*lyrata* Wachau tetraploids / *Lwt* hereafter; PIL, SCB, KAG, SWA, LOI, MAU). We found an additional tetraploid *lyrata* population in Hungary (GYE) and included it in this study. Diploid *lyrata* populations were chosen from the Wienerwald (PEQ, PER, VLH) which is the geographically closest diploid population to the *Let* and *Lwt*, and therefore a likely serve as source population.

For *arenosa*, representative populations of tetraploids from the Hercynian (WEK, SEN, BRD) and Alpine lineages (HOC, GUL, BGS) were selected as well as additional *arenosa* populations from the Western Carpathians (diploid: SNO; tetraploid: TRE), which is the centre of *arenosa* genetic diversity and the region of origin of the tetraploid cytotype^9^. For breadth, we selected several more diploid *arenosa* populations from the Pannonian (KZL, SZI) and Dinaric (BEL) lineages as well as the following populations from the hybrid zone in the eastern Austrian Forealps: HAL, ROK, FRE, OCH, and KEH. To complement our sampling with diploid *lyrata* from across its entire distribution range, we selected samples from the Hercynian (SRR2040791, SRR2040804), arctic-Eurasian (SRR2040796, SRR2040798, SRR2040805) and arctic-North American lineages (DRR054584, SRR2040769, SRR2040770, SRR2040789). *Arabidopsis croatica* (CRO) and *Arabidopsis halleri* (SRR2040780, SRR2040782, SRR2040783, SRR2040784, SRR2040785, SRR2040786, SRR2040787) were included as outgroups^37^. The majority of *lyrata* and hybrid samples were collected as seeds, cultivated and flash-frozen prior to DNA extraction, while samples for three populations (LIC, MOD, HAL) were collected and silica-dried. Silica-dried material from GYE was obtained from Marek Šlenker and Karol Marhold. *Arenosa* samples were collected and sequenced as part of a different study^21^. In addition, 16 accessions were downloaded from the NCBI Sequence Read Archive (SRA), bringing the total sample number to 92 (*SI Appendix*, Table S1). DNA of the *lyrata* and hybrid samples was extracted and purified from frozen or silica-dried leaf and/or flower tissue using the Epicentre MasterPure DNA extraction kit. DNA concentration measurements were performed with the Qubit 3.0 fluorometer (Invitrogen/Life Technologies, Carlsbad, California, USA). Genomic libraries for sequencing were prepared using the Illumina TRUSeq PCR-free library kit with 500 ng to 1 µg extracted DNA as input. We multiplexed libraries based on the Qubit concentrations, and those multiplexed-mixes were run on an initial quantification lane. According to the yields for each sample, loading of the same multiplex-mix on several lanes was increased to achieve a minimum of 15x coverage. Samples that had less than our target coverage were remixed and run on additional lanes. Libraries were sequenced as 125 bp paired-end reads on a HiSeq2000 by the Harvard University Bauer Core Facility (Cambridge, Massachusetts, USA).

### Data preparation and genotyping

Newly generated sequencing data and SRA accessions were processed together from raw fastq reads. We first used Cutadapt^43^ to identify and remove adapter sequences with a minimum read length of 25 bp and a maximum error rate of 0.15. We then quality trimmed reads using TRIMMOMATIC^44^ (LEADING:10 TRAILING:10 SLIDINGWINDOW:4:15 MINLEN:50). Samples sequenced on several lanes were then concatenated using custom scripts. Reads were deduplicated using ‘MarkDuplicates’ in picard v.1.103. Broadinst and readgroup names were adjusted utilizing ‘AddorReplaceReadGroups’ within the picard package. Reads were then mapped to the North American *lyrata* reference genome (v.2;^45^) using bwa-mem in the default paired-end mode^46^. Indels were realigned using the Genome Analysis Toolkit (GATK) ‘IndelRealigner’. Prior to variant discovery, we excluded individuals that had fewer than 40% of bases <8x coverage (assessed via GATK’s ‘DepthOfCoverage’ with the restriction to a minimum base quality of 25 and a minimum mapping quality of 25). Our final dataset for analysis contained 92 individuals.

Variant calling was performed using the GATK ‘HaplotypeCaller’ (--min_base_quality_score 25 --min_mapping_quality_score 25 -rf DuplicateRead -rf BadMate -rf BadCigar -ERC BP_RESOLUTION -variant_index_type LINEAR - variant_index_parameter 128000 --pcr_indel_model NONE), followed by ‘GenotypeGVCFs’ for genotyping. For each BAM file, ‘HaplotypeCaller’ was run in parallel for each scaffold with ploidy specified accordingly and retaining all sites (variant and non-variant). We combined the single-sample GVCF output from ‘HaplotypeCaller’ to multisample GVCFs and then ran ‘GenotypeGVCFs’ to jointly genotype these GVCFs, which greatly aids in distinguishing rare variants from sequencing errors. Using GATK’s ‘SelectVariants’, we first excluded all indel and mixed sites and restricted the remaining variant sites to be biallelic. Additional quality filtering was performed using the GATK ‘VariantFiltration’ tool (QD <2, MQ <40.00, FS >60.0, SOR >4.0, MQRankSum <-8.0, ReadPosRankSum <-8.0, DP <8). Then we masked sites that had excess read depth, which we defined as 1.6x the second mode (with the first mode being heterozygous deletions or mismapping) of the read depth distribution.

### Population structure

All analyses dedicated to reveal population structure and demography were based on putatively neutral 4-fold degenerate (4dg) SNPs only. We used the 4dg filter generated for *A. arenosa* from ^9^. After quality filtering these analyses were based on a genome-wide dataset consisting of 4,380,806 4dg SNPs, allowing for a maximum of 10% missing alleles per site (1.2% missing data) at a 5x coverage minimum for a given individual sample.

Although we expected fastSTRUCTURE^47^ to be superior in recognizing admixture compared to STRUCTURE^48^, running fastSTRUCTURE on our dataset resulted in poor performance, in that the result did not coincide with the STRUCTURE results or other analysis. This misbehaviour was probably due to the inclusion of polyploid data, as fastSTRUCTURE does not accommodate polyploid genotypes. We had randomly subsampled two alleles per each tetraploid site, similar to ^37^, using a custom script. But evidently such a subsampling strategy dissolves the fine-scale differences in admixture between populations at this scale. Hence, STRUCTURE was preferred, and was run on all samples and both ploidies. As STRUCTURE accepts only uniform ploidy as input, with one row per each ploidy, we added two rows of missing data for our diploid samples, making them pseudo-tetraploid. In addition, input data were linkage disequilibrium(LD)-pruned and singletons removed using custom scripts. Window size was set to 500 with a distance of 1000 between windows, allowing for 10% missing data, which resulted in a dataset of 32,256 SNPs genome-wide. We performed ten pruning replicates using the admixture model with uncorrelated allele frequencies, and then ran each for K values 2-10 with a burn-in period of 50,000 and 500,000 MCMC replicates. We conducted Principal Component Analysis (PCA) using the glPca function in the adegenet R package^49^.

### Demographic parameters and reconstruction of gene flow

We next performed demographic analyses with *fastsimcoal2*^23^ on 4dg sites. A minimum of two individuals per each population was required. Custom python scripts (FSC2input.py at https://github.com/pmonnahan/ScanTools/) were used to obtain the multi-dimensional allele frequency (DSFS) spectrum as well as bootstrap replicates of the DSFS for confidence interval estimation. For the bootstrap replicates, the genome was divided into 50 kb segments and segments were resampled with replacement until recreating a DSFS of equivalent size as the genome. Ultimately, we aimed to estimate demographic parameters and confidence intervals for a 4-population tree corresponding to diploid and tetraploid *lyrata* and *arenosa*. For computational efficiency, 3-population trees were initially used to establish the presence/absence of migration edges by comparing models with a single migration edge to a null model with no migration. Additional migration edges would then be added and compared to the initial simple model. For each model, 50 replicates were performed and values kept for the replicate with the highest likelihood. For each replicate, we allowed for 40 ECM cycles and 100,000 simulations in each step of each cycle for estimation of the expected side frequency spectrum (SFS). Although the above process identified the key migration edges, it resulted in a 4-population tree that was overly complex; the exercise suggested six migration edges in total (*SI Appendix*, Figure S3). Over-fitting was evidenced by highly imprecise and nonsensical estimates for a subset of parameters (*SI Appendix*, Table S2). For example, the ancestral population size for *lyrata* was estimated to be greater than 5 million with individual replicate estimates ranging from <100,000 to over 10 million. Estimates for population fusion times were also drastically greater than observed in previous 3-population trees. We therefore opted for a simpler model, retaining only the two migration edges with the highest support: bidirectional migration between tetraploids. Parameter estimates for each of the 100 bootstrap replicates were obtained using the scheme described above, and 95% confidence intervals were calculated using the 2.5^th^ and 97.5^th^ percentiles of the resulting distribution of each parameter.

### Changes in effective population size over time

Pairwise Sequentially Markovian Coalescent Model (PSMC) v.0.6.4 was used to infer changes in effective population size (*Ne*) through time using information from whole-genome sequences of *lyrata* and *arenosa* diploids^27^. We generated plots of the most deeply sequenced representative of each of the diploid *lyrata* and *arenosa* populations, with the exception of distinct *arenosa* KZL and SZI. A consensus FASTQ sequence was created using samtools v.1.2 and bcftools v.1.2 using samtools mpileup -C50 -Q 30 -q 30 with the *lyrata* v.2 genome as the reference. The reference was masked at all sites at which read depth was more than twice the average read depth across the genome. Samtools mpileup was followed with bcftools call -c and vcfutils.pl vcf2fq -d 5 -D 34 -Q 30 to create a FASTQ reference file. Using PSMC, this was changed to a format that was required with PSMC by fq2psmcfa -q20, and psmc was run with parameters psmc -N25 -t15 -r5 -p “4+25*2+4+6” and psmc_plot.pl –R -g 2 -u 3.7e-8 to get a text file that could be plotted with R. We used the mutation rate estimate µ=3.7×10^−8 9^ and a generation time of two years for both species, as *arenosa* is mainly biennial, and we estimate that *lyrata* generates the highest number of propagules in its second year after germination (R.S., personal observation).

### Introgression estimates using *D*_*FOIL*_

To test for introgression between populations of *lyrata* and *arenosa* in windows along the genome, we used the *D*FOIL approach^28^. *D*FOIL is an extension of the *D* statistic, or ABBA-BABA test^50^. It differs from the original *D* statistic in that it detects directional introgression in a 5-taxon phylogeny by using a combination of four separate *D* statistics, and by borrowing the logic of the “FOIL” (“First, Outer, Inner, Last”) method for multiplying two binomials in elementary algebra. We tested multiple 5-taxa scenarios on various tetraploid populations of *lyrata* and *arenosa*, using an *A. halleri* population as the outgroup in every scenario. *D*FOIL requires the first pair of taxa to have diverged prior to the second pair of taxa, so in each scenario the first pair of taxa included two populations of *arenosa*, and the second pair of taxa included two populations of *lyrata*.

### Cytological assessment of meiotic stability

Individual tetraploid *lyrata* and *arenosa* plants were germinated in 7 cm pots with Levington® Advance Seed and Modular Compost Plus Sand soil with 16h light / 8h dark cycles at 20°C constant temperature. Once rosettes had formed, plants were vernalised for six weeks with 8h light (6°C) / 16h dark (4°C) cycles. Plants were then grown in 16h light (13°C) / 8h dark (6°C) cycles to encourage flowering. Buds were collected, fixed and anthers dissected for basic cytology as described in ^51^ except that 50 mg (30 Gelatine Digestive Units) Zygest® Bromelain were added to the enzyme mixture, and incubation time was increased to 75 minutes. The prepared slides were stained and mounted with 7 µl DAPI in Vectashield (Vector Lab) and metaphase I chromosomes visualised using a Nikon 90i Eclipse fluorescent microscope with NIS Elements software. Chromosome spreads with all rod and/or ring bivalents were scored as “stable meiosis” (Figure 1D), while multivalents with multiple chiasmata were scored as “unstable meiosis” (Figure 1E).

### Differentiation scans for signatures of selective sweeps

We grouped populations by ploidy level, species or hybrid affiliation, and affiliation to a biogeographic region in case of tetraploid *lyrata*. We calculated the following metrics in adjacent nonoverlapping genomic windows: allele frequency differences (AFD), *dXY*, Fst^52^, Rho^53^, and the number of fixed differences between the *lyrata* diploids and the two *lyrata* tetraploid groups (*Let* and *Lwt*). We identified selective sweep candidates as the 1% outliers of the empirical distribution for each metric. To maximise our chances of capturing differentiation truly related to ploidy and not local adaptation, we selected the overlap between these two independent scans wherein the tetraploids contrast by edaphic (soil) preference and then focused on outliers were identified in a highly stringent genome scanning approach in *A. arenosa*^13^.

To obtain insight into differentiation between population groups, AFD, *dXY*, Fst, Rho, and the number of fixed differences were calculated for additional populations. *Arabidopsis arenosa* populations were grouped by lineage, as identified in ^9,21^, as ‘*arenosa* Hercynian tetraploids’ (*Aht*) and ‘*arenosa* Alpine tetraploids’ (*Aat*), which also corresponds to biogeographic groupings.

### Visualisation of allele frequencies

We visualized allele frequencies of amino acid substitutions in form of a heatmap. Pre-processed VCF files were annotated using SnpEff^54^ (https://dx.doi.org/10.4161%2Ff) with the manually added *lyrata* v.2 reference annotation^55^ (https://doi.org/10.1371/journa). Variants annotated as “missense” (i.e. amino acid substitutions) were extracted using SnpSift^54^. Gene coding loci were extracted from the whole-genome annotated VCF and per-population allele frequencies for each amino acid substitution calculated using GATK’s ‘SelectVariants’. Alternative allele frequencies (polarized against the *lyrata* reference) were visualized using the heatmap.2 function in the gplots package in R (Warnes et al., 2016, https://CRAN.R-project.org/pac).

### Identification of differentiated, introgressed regions

To investigate how the relationships among diploid and tetraploid populations of the two species vary across the genome, we used topology weighting by iterative sampling of subtrees (*Twisst*)^38^ (github.com/simonhmartin/twisst/). *Twisst* provides a quantitative measure of the relationships among a set of taxa when each taxon is represented by multiple individuals and the taxa are not necessarily reciprocally monophyletic. This provides a naive means to detect both introgression and incomplete lineage sorting, and how these vary across the genome. We first inferred genealogies for 50 SNP windows across the whole genome using the BIONJ algorithm^56^ as implemented in PHYML^57^. Because each individual carries two (for diploids) or four (for tetraploids) distinct haplotypes that represent different tips in the genealogy, it is necessary to first separate the haplotypes by phasing heterozygous genotypes. We used a heuristic approach to estimate phase that maximises the average extent of LD among all pairs of polymorphic sites in the window. This approach iteratively selects the best genotype configuration for each site, beginning with the site that has the most heterozygous genotypes. At each step, the optimal configuration is that which maximises the average LD between the target site and all previous target sites. This allows simultaneous phasing of diploids and tetraploids. We investigated the accuracy of this phasing approach using simulated sequences generated using the coalescent simulator *msms*^58^ and *seq-gen*^59^, following^38^, but here adding steps to randomise phase and then apply phase inference. Because *Twisst* is robust to within-taxon phasing errors^38^, the relevant question here is the extent to which imperfect phasing would affect the estimated topology weightings. We therefore applied *Twisst* to the simulated data and compared the results with (i) perfect phase, (ii) randomised phase, and (iii) randomised and then inferred phase. This confirmed that our heuristic phasing algorithm led to an improvement in the accuracy of the weightings, giving results that approached what is achieved with perfect phase information.

For running *Twisst* on the empirical data, we combined samples into four ingroup populations: diploid *lyrata*, tetraploid *lyrata*, diploid *arenosa* and tetraploid *arenosa*, and included *A. halleri* as an outgroup. These five taxa give fifteen possible rooted taxon topologies (Figure 3A, B). Although *Twisst* does not consider rooting when computing topology weightings, the inclusion of an outgroup improves the interpretation of the results, allowing the direction of introgression to be inferred in some cases^38^. In all analyses, topology weightings were computed exactly for all window trees that could be simplified to ≤2,000 remaining haplotype combinations (see ^38^ for details). In cases where this was not possible, approximate weightings were computed by randomly sampling combinations of haplotypes until the 95% binomial confidence interval for all fifteen topology weightings was below 0.05. Confidence intervals were computed using the Wilson method implemented in the package binom in R (R Core Team 2015).

## Data Availability

All sequence data are freely available in the NCBI Sequence Read Archive (SRA; https://www.ncbi.nlm.nih.gov/sra) with the primary accession code PRJNAXXXXX (available upon publication).

### Acknowledgements

This work was supported by the European Research Council (ERC) under the European Union’s Horizon 2020 research and innovation programme [grant number ERC-StG 679056 HOTSPOT], via a grant to L.Y.; and the Biotechnology and Biological Sciences Research Council [grant number BB/P013511/1], via a grant to the John Innes Centre (L.Y.). JDH was funded via BBSRC New Investigator grant BB/M01973X/1. Additional support was provided by the Charles University Grant Agency (GAUK 228716 to M.B.). Computational resources were partly provided by the CESNET LM2015042 and the CERIT Scientific Cloud LM2015085, under the programme “Projects of Large Research, Development, and Innovations Infrastructures”. The authors thank Jeff Doyle for critical reading of the manuscript.

## Author Contributions

LY and RS conceived the study. SM, PM, PJS, SHM, JK, PP, and MB performed analyses with input from LY, RS, and JDH. SM and PJS performed laboratory experiments. LY, RS and SM wrote the manuscript with input from all authors.

