## Supplement for "Interspecific introgression mediates adaptation to whole genome duplication"

### Supplementary Figures and Tables

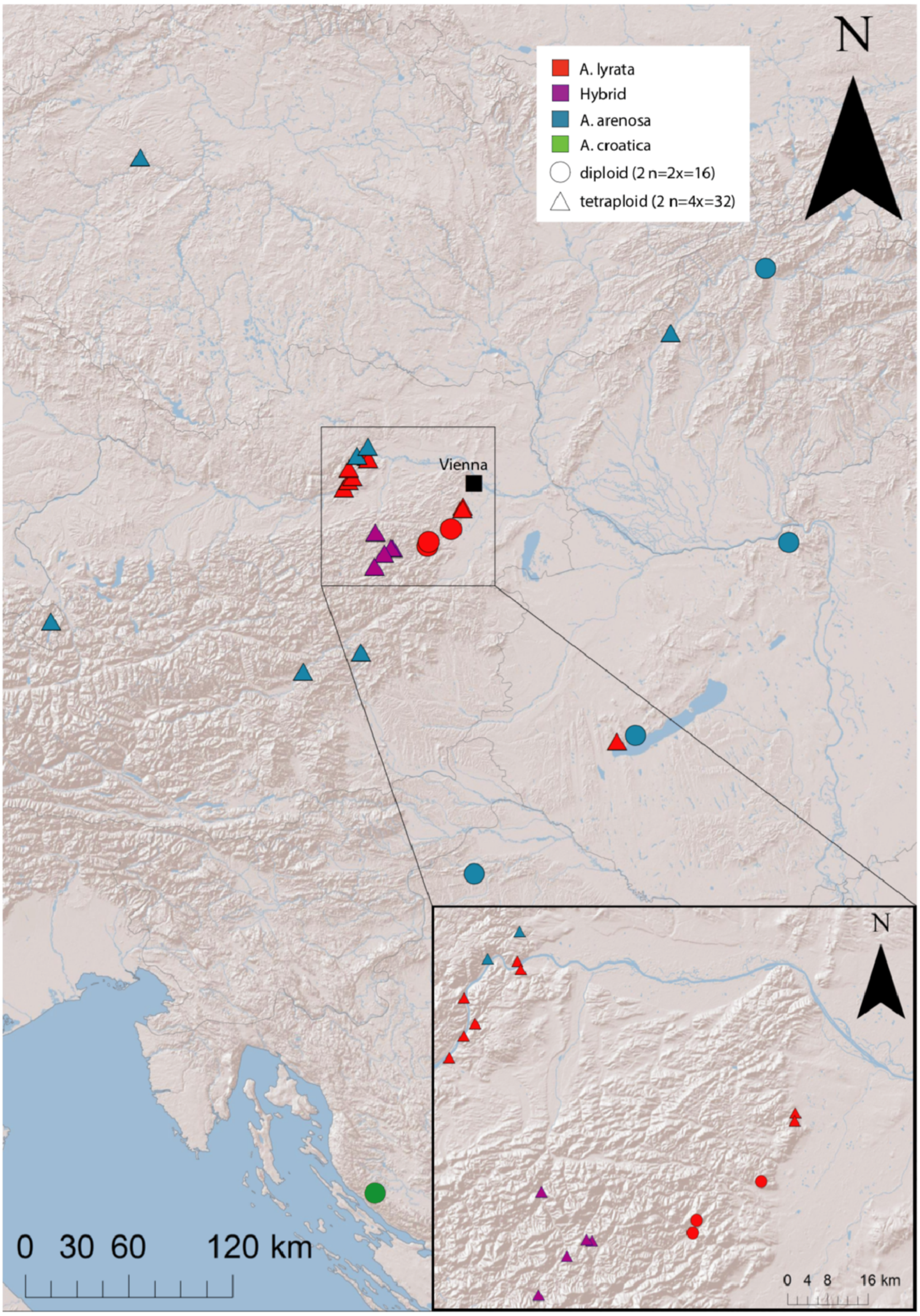


**Figure S1.** Map of Central Europe showing the locations for populations sampled in this study and from [21]. Circles represent diploid populations, triangles represent tetraploid populations. Colours are indicative of species and hybrids as indicated in the inset. The inset represents a zoomed in view of the eastern Austrian Forealps and the Wachau valley.

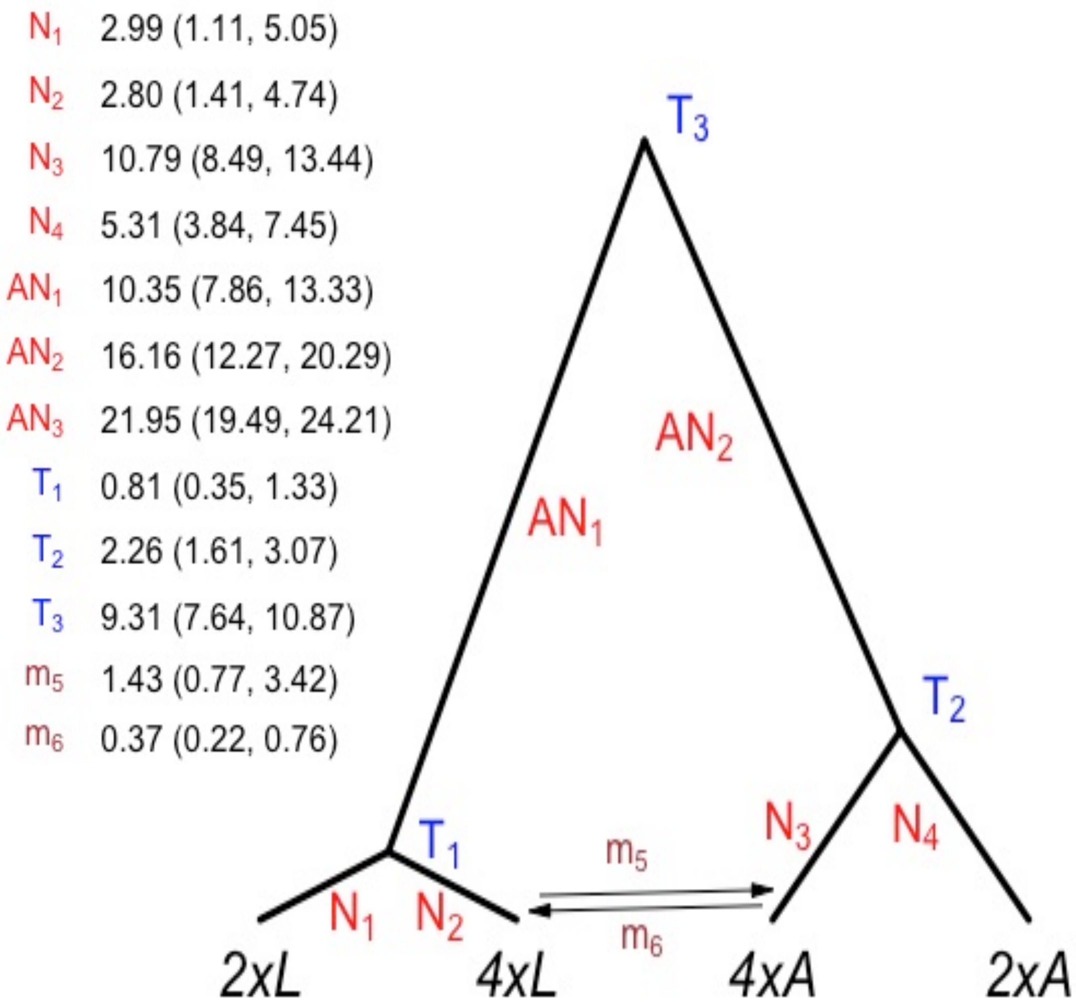


**Figure S2. Demographic parameter estimates for diploid *A. lyrata* (*2xL*), tetraploid *A. lyrata* (*4xL*), tetraploid *A. arenosa* (*4xA*), and diploid *A. arenosa* (*2xA*).** Units are 100,000's of individuals for population size (N and AN), 100,000's of generations for time estimates (T), and $10^{-6}$ alleles per generation. Median values across replicates are given.

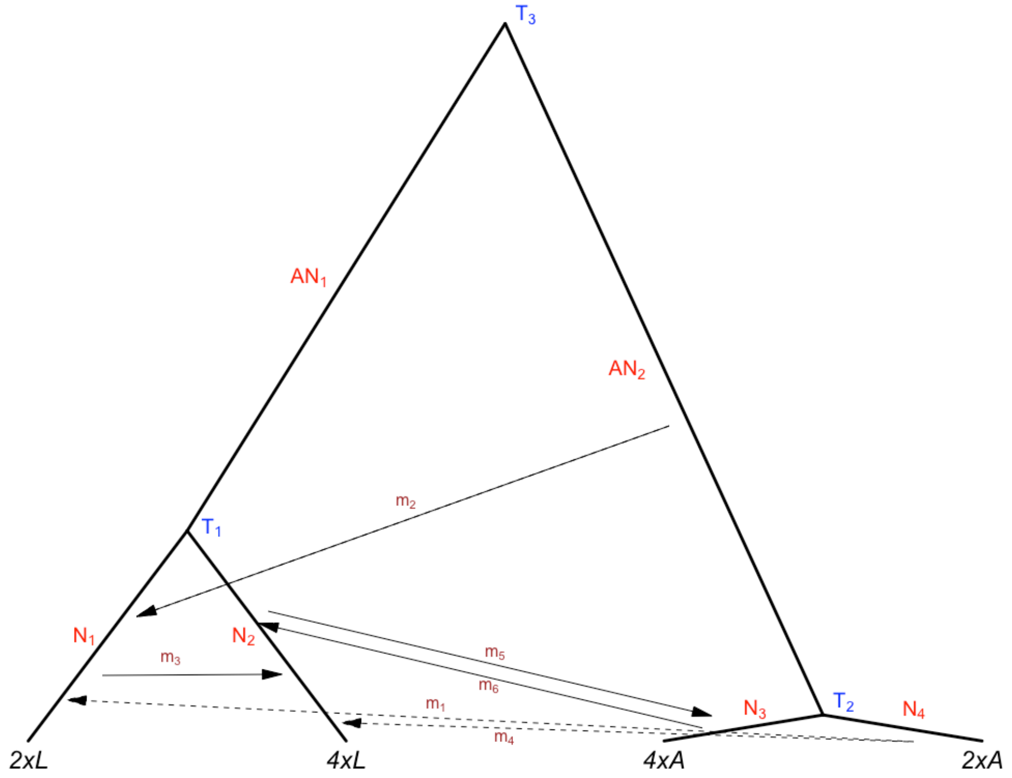


**Figure S3. Parameter estimates for the different 4 population scenarios**. Parameter values are given in Table S2.

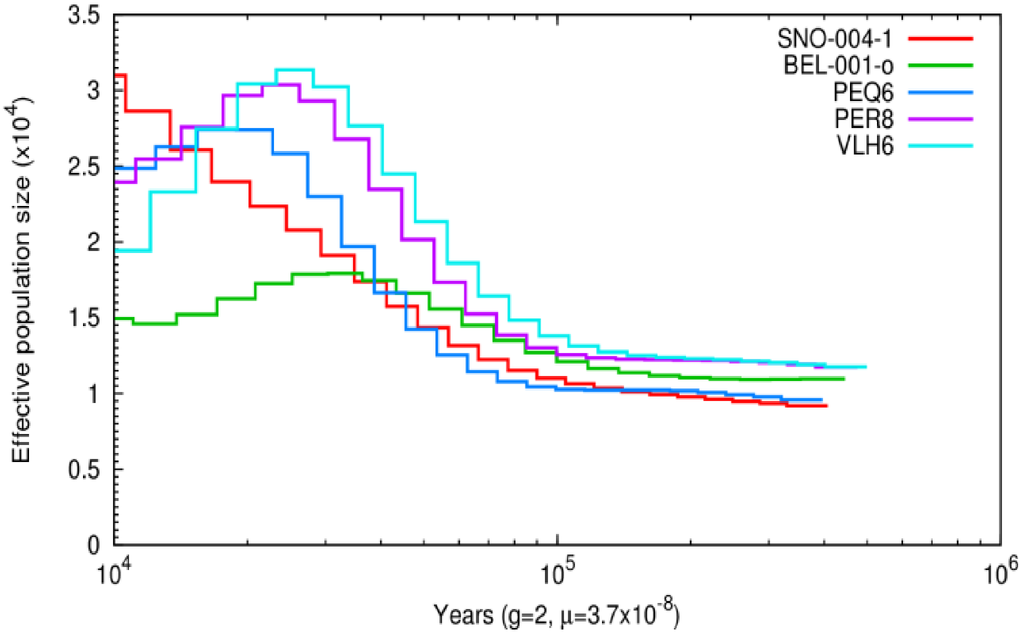


**Figure S4. Pairwise Sequentially Markovian Coalescent Model (PSMC) analysis of diploid *A. lyrata* and *A. arenosa* populations used in this study.** We used the mutation rate $\mu=3.7\times10^{-8}$ [9] and a generation time of two years for both species, as *A. arenosa* is mainly biennial, and we suppose that *A. lyrata* generates the highest number of propagules in its second year after germination.

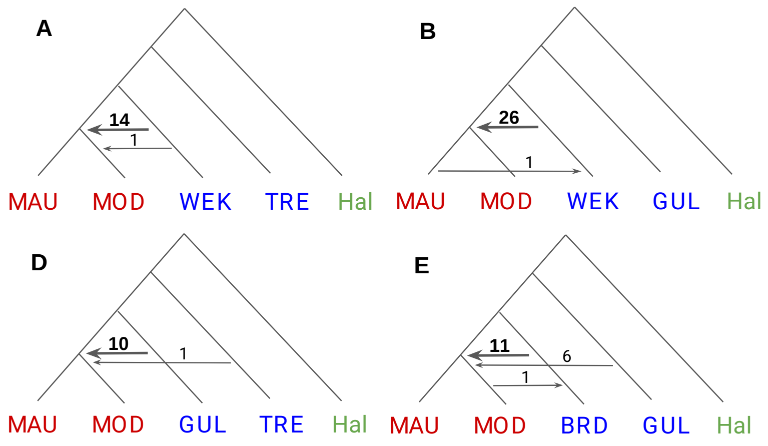


**Figure S5. Direction and frequency of introgression between pairs of autotetraploid *A. lyrata* and *A. arenosa* populations using $D_{FOIL}$ analysis.** Red: *A. lyrata*; blue: *A. arenosa*; green: *A. halleri*. Arrows show the direction of introgression. Numbers above the arrows indicate the number of introgression-indicative genomic windows.

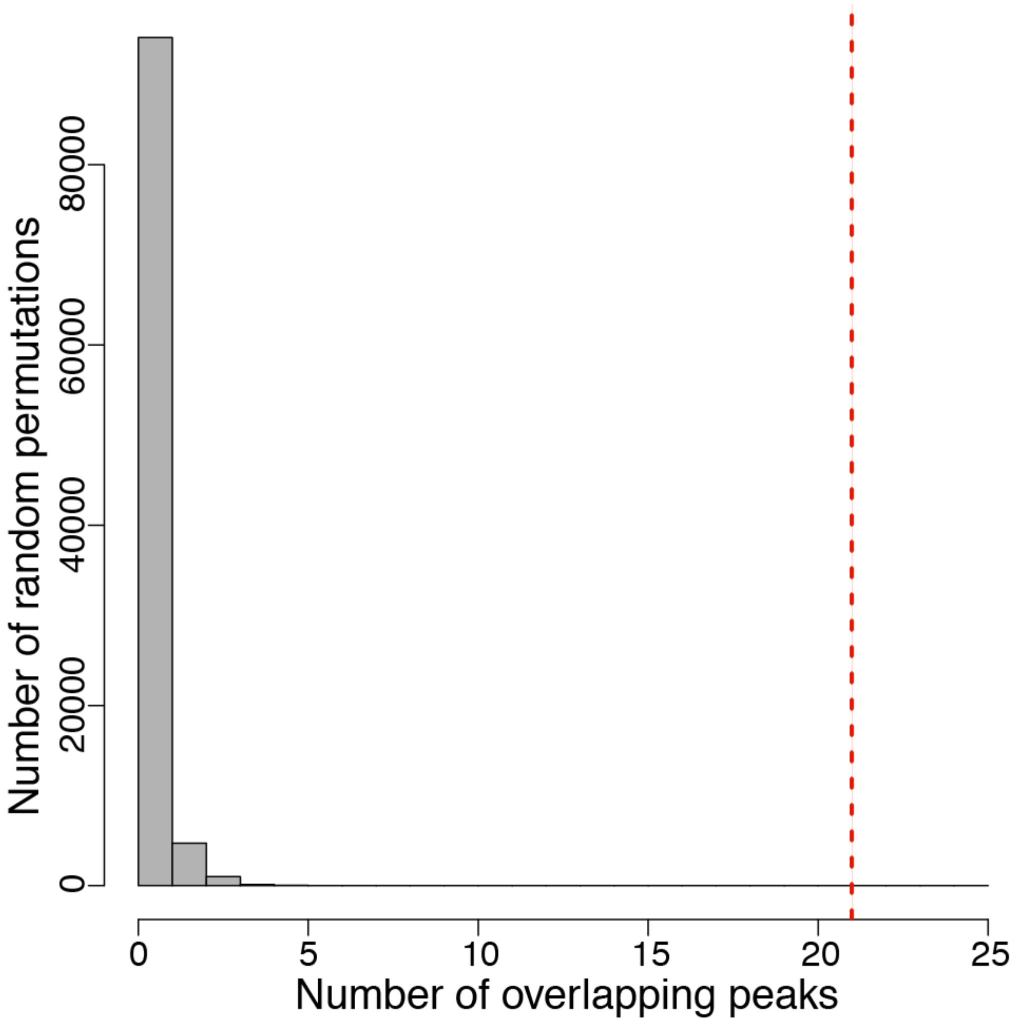


**Figure S6. Permutation tests indicating expected distribution of degree of overlap observed between *Twisst* outlier windows (61) and divergence scan outlier windows (195).** Red line indicates the observed value of 21 overlapping gene coding loci exhibiting both *Twisst* outlier and divergence scan outlier status, many more than expected by chance.

| Pop. | N ind. | Ploidy | Altitude | Latitude | Longitude | Country | Locality | Published in (orig. code) | Origin of tissue |
|---|---|---|---|---|---|---|---|---|---|
| BEL | 3 | 2x | 550 | 46.16167 | 16.11500 | HR | Castle ruin Belecgrad; open sites in the forest, walls of the castle ruin | Monnahan et al. 2019 (BEL) | wild |
| BGS | 3 | 4x | 570 | 47.62806 | 13.00167 | D | Berchtesgaden; railway, secondary gravel | Hollister et al. 2012 (BGS) | cultiv. |
| BRD | 2 | 4x | 350 | 50.04967 | 13.89081 | CZ | Brdatka; open forest in canyon of Berounka river | Monnahan et al. 2019 (BRD) | wild |
| CRO | 4 | 2x | 1076 | 44.53147 | 15.19402 | HR | Ljubičko Brdo and Zavižan; rocks | Monnahan et al. 2019 (CRO) | wild |
| FRE | 2 | 4x | 391 | 47.99405 | 15.57118 | AT | Freiland; rocks | | cultiv. |
| GUL | 3 | 4x | 820 | 47.29000 | 14.93167 | AT | Gulsen; serpentine rocks | Arnold et al. 2016 (GU) | cultiv. |
| GYE | 3 | 4x | 202 | 46.78442 | 17.28775 | HU | Gyenesdiás; dolomitic rock on northern border of the village | | wild |
| HAL | 3 | 4x | 665 | 47.90187 | 15.69233 | AT | Halbach valley; gravel | | wild |
| HOC | 3 | 4x | 580 | 47.37000 | 15.38667 | AT | Hochlantsch | Arnold et al. 2016 (HO) | cultiv. |
| KAG | 3 | 4x | 257 | 48.29432 | 15.42614 | AT | Wachau, Kartause Aggsbach; rocks | | cultiv. |
| KEH | 2 | 4x | 699 | 47.81611 | 15.54311 | AT | Kernhof; rocks, gravel | | cultiv. |
| KZL | 3 | 2x | 330 | 47.72444 | 18.77917 | HU | Kesztölc | Monnahan et al. 2019 (KZL) | cultiv. |
| LIC | 3 | 4x | 298 | 48.09283 | 16.27073 | AT | Liechtenstein castle; walls of the castle ruin, rocks | | wild |
| LOI | 3 | 4x | 289 | 48.39649 | 15.55268 | AT | Wachau, Loibenberg; oak-pine forest, rocks | | cultiv. |
| MAU | 3 | 4x | 244 | 48.3818 | 15.56031 | AT | Wachau, Mauternbach; oak-pine forest, rocks | | cultiv. |
| MOD | 3 | 4x | 335 | 48.07955 | 16.26718 | AT | Castle ruin Mödling; walls of the castle ruin, rocks | | wild |
| OCH | 2 | 4x | 698 | 47.87950 | 15.62691 | AT | Untermitterbach, road to Ochsattel; gravel | | cultiv. |
| PEQ | 2 | 2x | 461 | 47.90154 | 15.96864 | AT | Pernitz, small quarry in Haltergraben; rocks | | cultiv. |
| PER | 2 | 2x | 564 | 47.92251 | 15.98176 | AT | Pernitz, road from Pernitz to Pottenstein; rocks | | cultiv. |
| PIL | 3 | 4x | 224 | 48.23901 | 15.34931 | AT | Wachau, mouth of Pielach river into Danube river; rocks | | cultiv. |
| ROK | 2 | 4x | 662 | 47.90531 | 15.68304 | AT | Rossbachklamm; rocks, gravel | | cultiv. |
| SCB | 2 | 4x | 244 | 48.27428 | 15.39301 | AT | Wachau, Schönbühel; rocks | | cultiv. |
| SEN | 1 | 4x | 296 | 48.44750 | 15.56469 | AT | Castle ruin Senftenberg; walls of the castle ruin, rocks | | cultiv. |
| SNO | 2 | 2x | 390 | 49.17417 | 18.86167 | SK | Strečno | Yant et al. 2012 (SN) | cultiv. |
| SWA | 2 | 4x | 264 | 48.34031 | 15.40085 | AT | Wachau, Schwallenbach; rocks | | cultiv. |
| SZI | 3 | 2x | 130 | 46.80667 | 17.43444 | HU | Szigligeti vár | Monnahan et al. 2019 (SZI) | cultiv. |
| TBG | | 4x | 640 | 48.13972 | 8.23667 | D | Triberg; railway, secondary gravel | Hollister et al. 2012 (TBG) | cultiv. |
| TRE | 3 | 4x | 280 | 48.89417 | 18.04472 | SK | Trenčín; rocks at the castle ruin | Monnahan et al. 2019 (TRE) | cultiv. |
| VLH | 3 | 2x | 484 | 47.97978 | 16.16374 | AT | Vöslauer Hütte; pine forest, rocks | | cultiv. |
| WEK | 3 | 4x | 359 | 48.40502 | 15.47291 | AT | Wachau, Weißenkirchen; oak-pine forest, former vineyard | Monnahan et al. 2019 (WEK) | wild |

**Table S1. The 30 populations included in this study.**

| Variable | Model1 | Model2 | Model3 | Model4 | Model5 | Model6 |
|---|---|---|---|---|---|---|
| N1 | 465938 | 482982 | 437237 | 303955 | 110886 | 369141 |
| N2 | 75834 | 88151 | 188323 | 289967 | 373998 | 339673 |
| N3 | 836187 | 873906 | 927130 | 1079527 | 1120090 | 1091553 |
| N4 | 561181 | 461999 | 474882 | 540394 | 566028 | 473956 |
| AN1 | 4959828 | 4376433 | 3004479 | 1036598 | 1162941 | 934617 |
| AN2 | 2036501 | 2011382 | 1901016 | 1611485 | 1169502 | 1746457 |
| AN3 | 2574717 | 2003004 | 2072445 | 2207793 | 2354880 | 2249547 |
| T1 | 2034971 | 383815 | 253022 | 81326 | 47876 | 99582 |
| T2 | 217193 | 182807 | 190528 | 230060 | 247875 | 210015 |
| T3 | 6589497 | 1238849 | 1095980 | 930721 | 613422 | 838791 |
| m1 | 2.79E-07 | -- | -- | -- | -- | -- |
| m2 | 7.48E-07 | -- | -- | -- | | -- |
| m3 | 1.71E-05 | 2.44E-05 | 8.28E-06 | -- | -- | -- |
| m4 | 1.18E-06 | 1.28E-06 | -- | -- | -- | -- |
| m5 | 3.61E-07 | 5.80E-07 | 4.30E-07 | 4.29E-07 | 8.14E-07 | -- |
| m6 | 2.71E-06 | 4.34E-06 | 2.21E-06 | 1.57E-06 | -- | 1.26E-06 |
| Likelihood | -623798 | -622710 | -622579 | -622342 | -625995 | -622483 |
| AIC | 2872730 | 2867798 | 2867139 | 2865432 | 2882840 | 2866662 |

**Table S2. Parameter estimates, likelihood, and AIC for different 4-population scenarios in *fastsimcoal2*.** Model 4 with the highest likelihood and lowest AIC was chosen. Mean values are given, in contrast to medians in Figure 1C and Figure S2.

| Population | Plant no. | No. cells with MIs scored | No. cells with stable MIs | No. cells with unstable MIs | % stable | % unstable |
|---|---|---|---|---|---|---|
| **LIC** | 1 | 13 | 13 | 0 | 100 | 0 |
| | 2 | 14 | 3 | 11 | 21 | 79 |
| | 3 | 21 | 4 | 17 | 19 | 81 |
| | 4 | 49 | 40 | 9 | 82 | 18 |
| | 5 | 22 | 0 | 22 | 0 | 100 |
| | 6 | 13 | 10 | 3 | 77 | 23 |
| **MOD** | 1 | 20 | 13 | 7 | 65 | 35 |
| | 2 | 67 | 58 | 9 | 87 | 13 |
| | 3 | 8 | 7 | 1 | 88 | 13 |
| | 4 | 25 | 1 | 24 | 4 | 96 |
| **KAG** | 1 | 23 | 21 | 2 | 91 | 9 |
| | 2 | 35 | 34 | 1 | 97 | 3 |
| | 3 | 30 | 0 | 30 | 0 | 100 |
| | 4 | 42 | 29 | 13 | 69 | 31 |
| | 5 | 39 | 27 | 12 | 69 | 31 |
| **ROK** | 1 | 17 | 0 | 17 | 0 | 100 |
| | 2 | 38 | 30 | 8 | 79 | 21 |
| | 3 | 25 | 10 | 15 | 40 | 60 |
| | 4 | 37 | 10 | 27 | 27 | 73 |
| | 5 | 47 | 13 | 34 | 28 | 72 |
| **WEK** | 1 | 18 | 2 | 16 | 11 | 89 |
| | 2 | 30 | 29 | 1 | 97 | 3 |
| | 3 | 15 | 13 | 2 | 87 | 13 |
| | 4 | 26 | 26 | 0 | 100 | 0 |
| | 5 | 35 | 30 | 5 | 86 | 14 |
| **SEN** | 1 | 25 | 0 | 25 | 0 | 100 |
| | 2 | 29 | 2 | 27 | 7 | 93 |
| | 3 | 22 | 2 | 20 | 9 | 91 |
| | 4 | 29 | 2 | 27 | 7 | 93 |
| | 5 | 16 | 3 | 13 | 19 | 81 |
| **TBG** | 1 | 47 | 32 | 15 | 68 | 32 |
| | 2 | 29 | 3 | 26 | 10 | 90 |
| | 3 | 52 | 37 | 15 | 71 | 29 |
| | 4 | 22 | 6 | 16 | 27 | 73 |
| | 5 | 13 | 9 | 4 | 69 | 31 |

**Table S3. Chromosome stability scoring of individual plants from tetraploid populations of *A. lyrata* and *A. arenosa* and a hybrid population at meiotic metaphase I (MI).** Chromosome spreads with all rod and/or ring bivalents were scored as “Stable meiosis” (Figure 1D), while multivalents with multiple chiasmata were scored as “Unstable meiosis” (Figure 1E). Tetraploid *A. lyrata*: LIC, MOD, KAG. Tetraploid *A. arenosa*: WEK, SEN, TBG. The TBG population was not integrated in the other parts of this study, but is included for comparison; it was the tetraploid *A. arenosa* population on which the study of [13] was based.

| | Contrast | No. SNPs | AFD | $d_{XY}$ | Fst | Rho | Fixed Diff |
|---|---|---|---|---|---|---|---|
| *Lyrata* diploid vs. tetraploid | *Lyd* vs. *Let* | 2,904,110 | 0.14 | 0.22 | 0.09 | 0.19 | 270 |
| | *Lyd* vs. *Lwt* | 3,794,257 | 0.11 | 0.16 | 0.07 | 0.17 | 64 |
| *Lyrata* tetraploid vs. tetraploid | *Let* vs. *Lwt* | 4,795,381 | 0.09 | 0.16 | 0.06 | 0.13 | 24 |
| *Arenosa* tetraploid vs. tetraploid | *Aht* vs. *Aat* | 1,812,223 | 0.10 | 0.16 | 0.03 | 0.07 | 0 |
| *Lyrata* vs. *arenosa* | *Lyd* vs. *Aht* | 1,729,114 | 0.25 | 0.27 | 0.39 | 0.39 | 41,810 |
| | *Lyd* vs. *Aat* | 2,874,610 | 0.23 | 0.24 | 0.40 | 0.40 | 57,492 |
| | *Let* vs. *Aht* | 2,257,560 | 0.21 | 0.24 | 0.34 | 0.36 | 17,000 |
| | *Lwt* vs. *Aht* | 2,513,764 | 0.15 | 0.19 | 0.26 | 0.32 | 767 |
| | *Let* vs. *Aat* | 3,644,666 | 0.20 | 0.23 | 0.35 | 0.37 | 21,372 |
| | *Lwt* vs. *Aat* | 3,653,076 | 0.16 | 0.20 | 0.29 | 0.34 | 947 |
| Hybrids from the eastern Austrian Forealps vs. *lyrata* or *arenosa* | *Hy1* vs. *Lyd* | 4,165,349 | 0.16 | 0.20 | 0.17 | 0.29 | 259 |
| | *Hy2* vs. *Lyd* | 4,055,168 | 0.16 | 0.20 | 0.15 | 0.27 | 322 |
| | *Hy1* vs. *Aat* | 3,797,036 | 0.11 | 0.18 | 0.11 | 0.20 | 2 |
| | *Hy2* vs. *Aat* | 3,752,996 | 0.13 | 0.19 | 0.15 | 0.24 | 43 |

**Table S4. Differentiation between various contrasts.** Genome-wide metrics of differentiation are allele frequency differences (AFD), $d_{XY}$, Fst, Rho, and the number of fixed differences (Fixed Diff). Diploid *A. lyrata*: *Lyd*. Tetraploid *A. lyrata*: *lyrata* eastern tetraploids (*Let*), *lyrata* Wachau tetraploids (*Lwt*). Tetraploid *A. arenosa*: *arenosa* Hercynian tetraploids (*Aht*), *arenosa* Alpine tetraploids (*Aat*). Hybrids from the eastern Austrian Forealps: HAL, ROK, FRE, OCH, KEH (*Hy1*), and, alternatively, HAL, ROK, FRE, OCH (*Hy2*); the distinction between *Hy1* and *Hy2* was made because only HAL, ROK, FRE, and OCH are intermediate hybrids (KEH is more *arenosa*-like).

### Supplementary Datasets:

**Supplementary Dataset S1**: List of the top 1% outliers from the genome scan of diploid *A. lyrata* vs. *Let* ("*Let* scan") and diploid *A. lyrata* vs. *Lwt* ("*Lwt* scan"). The overlapping outlier loci are shown in a separate tab.

**Supplementary Dataset S2.** Genomic windows with weightings above 0.5 for topologies 6, 11 and 14 in the *Twisst* analyses. Genes-coding loci found in both *Twisst* analyses are indicated in bold.
